# Disentangling Prediction and Feedback in Social Brain Networks: A Predictive Processing Approach

**DOI:** 10.1101/2025.06.10.658968

**Authors:** Yiyu Wang, Juliet Y. Davidow, Richard D. Lane, Ajay B. Satpute

## Abstract

While much research in social cognitive neuroscience has focused on which brain regions are engaged when processing social content, it remains unclear what these areas are doing in terms of underlying mechanisms. Here, we approached this question using predictive processing theory, which suggests that the brain instantiates a generative model of its sensory environment. Using a novel animated shapes fMRI task, we observed a functional double dissociation between brain regions that were engaged when forming a prediction for agentic movement - which involved the premotor cortex and the lateral parietal cortex, previously implicated in action observation and mirroring - from those associated with processing feedback for updating abstract priors to guide predictions - which involved the dorsomedial and ventrolateral prefrontal cortex, the temporoparietal area, and the lateral temporal cortex, previously associated with mentalizing/theory of mind. We observed parallel functional dissociations in the cerebellar areas affiliated with these networks. These findings suggest new insights into how brain regions associated with action observation/mirroring and mentalizing/theory of mind play complementary roles in supporting facets of a predictive processing architecture underlying social cognition.

## Introduction

According to predictive processing theories, the brain is thought to predict its sensory environment by creating internal models that anticipate incoming input and using feedback to update those models when expectations are violated (Clark, 2013; Friston, 2010; Rao & Ballard, 1999). Higher-level cortical regions issue predictions that cascade down a neural hierarchy, while deviations from predictions are propagated back up the hierarchy to update the generative model (Clark, 2013; Rao & Ballard, 1999)(Figure 1). The predictive processing architecture offers a metabolically efficient and integrative computational account of neural information processing (Bastos et al., 2012; Spratling, 2017) that has been particularly influential in explaining sensory and motor processing (Barrett & Simmons, 2015; Clark, 2013; Heilbron & Chait, 2018; Rao & Ballard, 1999; Seth & Friston, 2016). Yet, its implications for understanding social cognition (Brown & Brüne, 2012; S. Gallagher & Allen, 2018; Keysers et al., 2024; Westra, 2019), among other complex psychological domains (Alexander & Brown, 2018; Clausi et al., 2019; Lee et al., 2021; Lewis & Bastiaansen, 2015; Satpute et al., 2020), remain comparatively underexplored. This gap is particularly significant, as social behavior poses one of the most complex prediction challenges for the brain to deal with (Ho et al., 2022; Tamir & Thornton, 2018). Other people act autonomously and dynamically, introducing considerable variability into the sensory stream. Theoretical accounts suggest that individuals manage this uncertainty by forming “mental models” (Ho et al., 2022; Koster-Hale & Saxe, 2013; Tamir & Thornton, 2018; Van Overwalle et al., 2020)— which for predictive processing models refer to priors that forecast how social behavior relates to changes in our sensory environment— to anticipate others’ actions, and by processing feedback to revise those models (Atzil et al., 2018; Brown & Brüne, 2012; Frith & Frith, 2006; S. Gallagher & Allen, 2018; Gendron & Barrett, 2018; Keysers et al., 2024).

**Figure 1.**
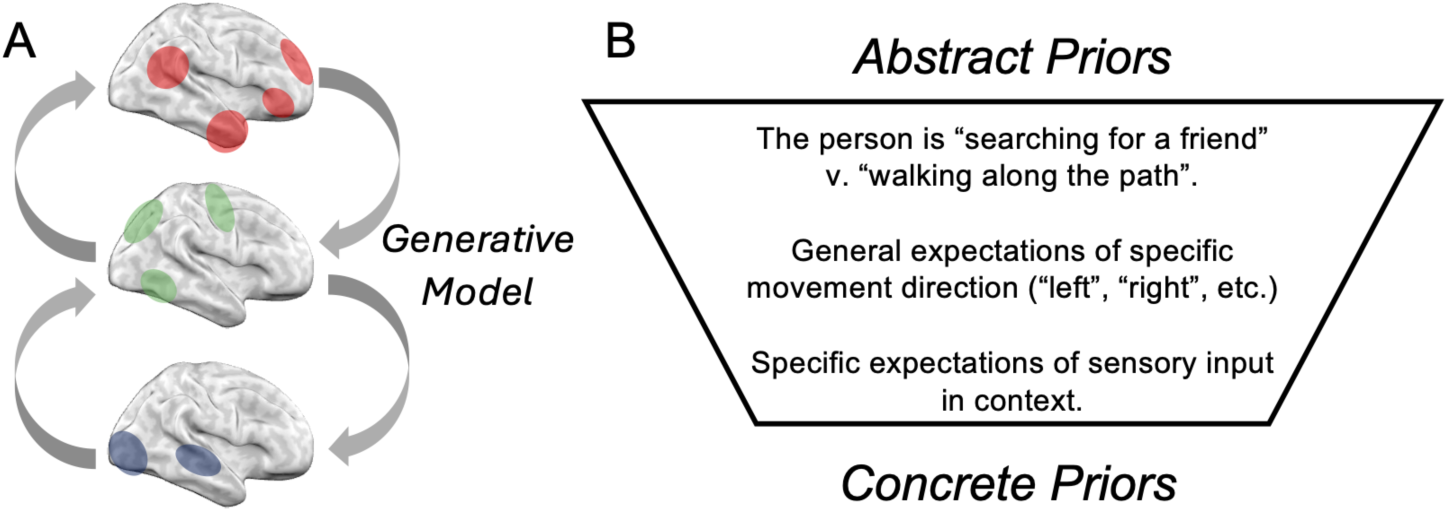
Social prediction and agentic movement perception. Social agents introduce considerable variation in the sensory environment. To navigate the social world, people flexibly deploy mental models to make predictions of others’ movements. In the cartoon scene, future movements may be predicted using a combination of relatively more abstract priors (e.g., people tend to walk in a straight line; people interacting with each other will move in ways that are contingent on each other, etc.), and increasingly more concrete priors (the general direction that the person will move next, or concrete sensory expectations of visual input). (A) According to predictive processing theories, the brain instantiates a generative model that predicts sensory input and processes perturbations as feedback to update the generative model. The figure shows a schematic of this model wherein areas distant from early exteroceptive sensory input (top, in red) represent more abstract priors and send cascading predictions down the hierarchy, while those lower in the hierarchy represent increasingly more concrete priors (at the bottom) and send prediction errors up the hierarchy. Deploying and updating priors with feedback may occur at many different levels of the hierarchy, depending on the nature of the information in question. (B) On a psychological level, we refer to the brain’s generative model as a “mental model”, which refers to a set of information processing priors that vary in abstraction. In the present study, we examine neural correlates associated with prediction generation during the cue period, and model updating associated with relatively abstract, “agent-dependent” and “agent-independent” priors.

Here, we investigated the neural mechanisms that support the flexible use and updating of mental models when predicting autonomous, agentic movements(Schultz & Frith, 2022). We reasoned that because social input is often ambiguous and context dependent (Barrett, 2022; Barrett et al., 2019; FeldmanHall & Shenhav, 2019; Heider & Simmel, 1944; Zaki & Ochsner, 2011), people must be able to learn and deploy different mental models based on feedback. Even in a relatively simple social environment—like predicting how people move in a public space —one might rely on different kinds of mental models. For example, a person could use a simple trajectory model (e.g., people tend to walk in a straight line) or a socially interactive model (e.g., people adjust their movement based on others), leading to different predictions from the same sensory input. Sensory feedback that is congruent or incongruent with predictions is then used to update the mental model.

To investigate the functional neural architecture underlying mental model formation and updating with feedback, we developed a novel task that separates these dynamic processes into distinct stages. Participants viewed cue videos showing two agents depicted as abstract shapes in a given setting, made a behavioral prediction of the movement of one of the agents, and were then shown a feedback video of where the agent ended up going. Each unique cue video could be interpreted using one of two abstract priors, one in which the target agent’s future behavior would ultimately depend on the other agent’s behavior (agent-dependent) or not (agent-independent), respectively, to form a mental model for making a unique prediction. We manipulated the proportion of feedback videos to be more consistent with one or the other prior such that participants were required to flexibly deploy different mental models to guide their predictions across trials. We validated the novel task paradigm behaviorally (N = 110), showing that participants adopted distinct mental models to guide their agentic movement predictions based on feedback. We then examined the functional neural correlates of these processes using fMRI (N = 30).

Based on past literature, we expected brain regions previously associated with mentalizing and theory of mind - including the anterior medial prefrontal cortex, precuneus, and lateral temporal cortex (Amodio & Frith, 2006; Bzdok et al., 2012; H. L. Gallagher & Frith, 2003; Satpute & Lieberman, 2006; Wheatley et al., 2007) - and/or those associated with action observation and action planning - including the premotor cortex, lateral parietal cortex, and lateral occipital cortex (Caspers et al., 2010; Decety & Grèzes, 2006; Iacoboni, 2009; Molenberghs et al., 2012) - to be associated with agentic movement perception (Spunt et al., 2011; Van Overwalle & Baetens, 2009). However, the nature of their involvement from a predictive processing view is unclear (see Cerliani et al., 2022; Keysers & Gazzola, 2014; Kilner et al., 2007). One hypothesis is that the same network involved in forming mental models that guide prediction also supports feedback processing for updating models; another is that these processing stages rely on distinct networks. A second set of hypotheses concerns the features of mental models being represented: agent-dependent and agent-independent priors may rely on overlapping or dissociable functional circuits. Finally, a third set of hypotheses concerns how prediction errors are represented. Incongruent feedback may engage a set of brain regions regardless of the mental model in use (condition-general prediction error), or alternatively, it may vary depending on the specific mental model being deployed (condition-dependent prediction error). We employed a data-driven analytical approach to test these possibilities. This approach made it possible to characterize dominant sources of variance in driving functional activity in each brain region. It further allowed us to evaluate whether the areas that are most influenced by the task are the same as those previously implicated in mentalizing and/or action observation and planning, and further, whether these areas cluster into previously established networks but on the basis of their task-related functional activation patterns.

## Methods

### Participants

A behavioral study involving consenting neurotypical adult college students at Northeastern University (N = 110; 70 females, 40 males, 0 non-identified/other; age range = [18, 22]) was conducted to validate the novel task. The fMRI study involved a separate sample of neurotypical participants between the ages of 18 to 40 years recruited from the greater Boston area and who received monetary compensation for participation. Participants were excluded if they reported positive indication of claustrophobia, a psychiatric diagnosis, use of psychotropic medication, being non-right-handed, having metal in the body, or if they exhibited excessive motion (N = 5; criteria described below). Participants completed an unrelated affective processing task before the present task. The effective fMRI sample for analysis included 30 participants (11 female, 19 male, 0 non-identified/other; age range [18, 38] years).

### Stimuli

Our stimulus set was inspired by the classic animated shapes task developed by Heider and Simmel (1944), in which participants watched a video of geometric shapes moving on a screen and were asked to describe what they saw. Despite the abstract nature of the stimuli, people often interpreted the movements using mental models (e.g., the agents were “chasing” each other or “hiding”). Variants of this task, including the Frith-Happé animations, have profoundly influenced social cognitive neuroscience, serving as the basis for early neuroimaging studies (e.g., Castelli et al., 2004; Martin & Weisberg, 2003) and as a task for the social cognition domain in the large-scale, Human Connectome Project (Van Essen et al., 2013). This paradigm revealed how observers impose animacy even upon abstract stimuli, offering a window into the mental models people use to make meaning from inherently ambiguous sensory input.

The goal of our unique stimulus set was to investigate functional neural activity associated with developing a mental model for forming predictions and processing feedback of agentic movement (cf. Cerliani et al., 2022). To achieve this, we created a novel stimulus set consisting of forty brief cue videos (4-8 seconds) with each one displaying a unique and stationary two-dimensional setting (e.g., a box, a cross, a set of dots) and the movement of two agents: a green square and a yellow square. For each cue video, we created two feedback videos (2-6 seconds), one in which the green square would follow or approach the yellow square (agent-dependent condition), and one in which the green square would continue a trajectory independently from the yellow square (agent-independent condition). Cue videos were designed such that participants would make different movement predictions depending on whether they were using agent-dependent or agent-independent priors.

### fMRI Task

The fMRI task and trial layout is presented in Figure 2. On each trial, participants saw a cue video (randomly chosen from the stimulus set) that paused on the final frame. They had four seconds to predict the green square’s movement using relative directions [left, right, up, or down]. After a jittered fixation interstimulus interval (1–2 seconds), the feedback video played. Participants then rated how surprised they were by the feedback using a sliding scale from “low” to “high” (4 seconds), followed by a jittered fixation intertrial interval (1–3 seconds).

**Figure 2.**
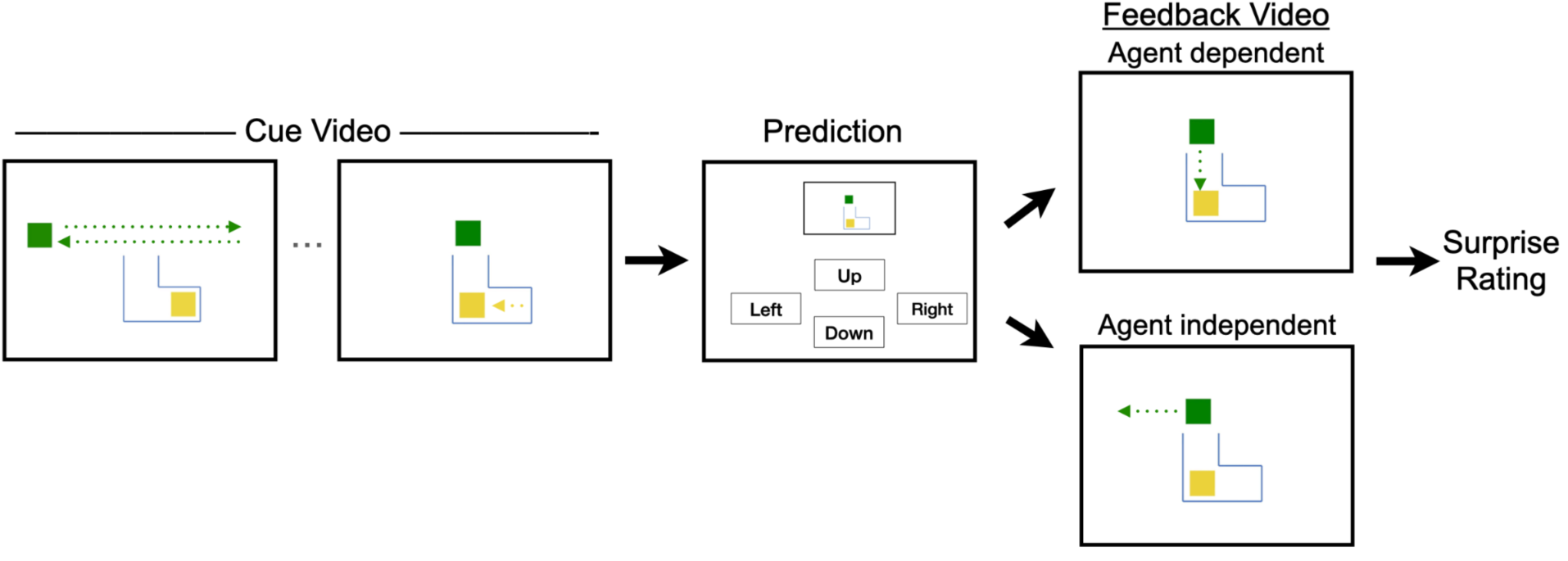
fMRI Task. Each trial began with a unique cue video (see Figure S1 for other examples; download the stimulus set at https://github.com/yiyuwang/SocialPrediction_AnimatedShapesTest). In this example, the green square moved back and forth along the dashed trajectory during which a yellow square moved “into view” for the green square. The video paused, participants entered their prediction for the green square’s movement (4 seconds), a feedback video was shown that depicts movement consistent with either agent dependent or agent independent priors, and then participants rated their surprise (4 seconds). Jittered interstimulus (1-2 seconds) and intertrial intervals (1-3 seconds) were included to separate brain imaging signals during cue and feedback videos within and across trials.

All behavioral responses were made using a trackball. We manipulated the proportion of agent-dependent or agent-independent feedback videos over trials to increase the utility of using one or the other mental model when viewing cues and making predictions on future trials. Specifically, trials were organized using a pseudo-probabilistic design (70/30 split) across two functional runs. In one run, the first five trials showed only agent-dependent feedback to set a prior early in the run; the subsequent fifteen trials were block randomized into three blocks consisting of three trials with agent-dependent feedback and two trials with agent-independent feedback. The other run was the same but favored agent-independent feedback. Condition order was counterbalanced across participants. Critically, the same exact cue videos could be followed by either type of feedback video. The feedback videos made it more optimal to conceptualize the cue video using agent-dependent or agent-independent priors to make more successful predictions.

### Behavioral Study

Prior to the fMRI study, a separate behavioral study was conducted to validate the novel stimuli and task paradigm. The behavioral study was used to examine how participants update their predictions with trial-by-trial feedback, and how they experienced engaging in the task. The design was similar to the fMRI study but included additional measures. After the cue video and prediction, participants rated their confidence in their prediction (“low” to “high”, [0-1], sliding scale) and then described the cue movement they viewed by typing in a text box. After the feedback video and the surprise rating, they again described the movement they viewed using a text box. Finally, a deterministic design was used to maximize the effects of feedback on future predictions. Participants were randomly assigned to receive only agent-dependent feedback for the first 20 trials followed by agent-independent feedback for the next 20 trials, or vice versa. The change in feedback condition occurred on trial 21 without disruption to the task or explicit indication to the participant. Text data per trial were analyzed using Linguistic Inquiry Word Count (LIWC-22; Boyd et al., 2022), from which we extracted content measures including use of personal and impersonal pronouns and use of social words, which we expected to differ by agent-dependent and agent-independent trial blocks.

### fMRI data acquisition and preprocessing

MRI data were collected using a 3T Siemens Prisma MRI scanner. Functional images were acquired in interleaved order using a T2*-weighted multiband echo planar imaging (EPI) pulse sequence (transverse slices, TR=800 ms, TE = 37 ms, flip angle=52°, FOV = 208mm, 2 mm thickness slices, voxel dimension= 2 x 2 x 2 mm, Phase Encoding Direction anterior to posterior (AP)). Anatomical images were acquired at the end of the session with a T1-weighted pulse sequence (TR = 2500ms, TE = 1.81ms, flip angle=8°, FOV = 256 mm, 0.8 mm thickness slices, voxel dimension= 0.8 x 0.8 x 0.8 mm, Phase Encoding Direction anterior to posterior (AP)). Image volumes were preprocessed using fMRIprep (Esteban et al., 2019). Preprocessing included motion correction, slice timing correction, removal of low frequency drifts using a temporal high-pass filter (discrete cosine transform, 0.01Hz cutoff), spatial smoothing (6mm FWHM). For all analyses, functional volumes were registered to participants’ anatomical image and then to a standard template (MNI152) using FSL flirt with boundary-based registration (BBR) metric - 6 dof (Jenkinson et al., 2002). Participants (N = 5) with head motion > 3 mm maximum framewise displacement, or a mean framewise displacement > 0.3, or > 5% of frames with motion spikes larger than 3mm in a single run, were excluded from analysis.

### Analysis

The task is designed such that the same cue videos can be flexibly interpreted using agent-dependent or agent-independent abstract priors, leading to distinct movement predictions on each trial. Thus, cue videos during each trial were assigned to mental model conditions based on participants’ predictions, and feedback videos were assigned to congruent and prediction error conditions based on whether the predictions were confirmed or disconfirmed, respectively.

A general linear model (GLM) was fitted to the fMRI data using Nilearn’s FirstLevelModel function. The GLM was fitted to the two runs concatenated together, and included a run regressor. The GLM included separate boxcar regressors for cue period, congruent feedback, and prediction error feedback, relative to agent-dependent and agent-independent responses (six condition regressors). The regressors were convolved with the canonical double gamma hemodynamic response function based on the run before the concatenation. Nuisance regressors included standard motion parameters (three for translation and three for rotation), physiological noise artifacts (one regressor each for cerebrospinal fluid, white matter, and framewise displacement from fMRIprep output), and non-steady state outliers (separate stick functions for each outlier volume). On average out of the 40 trials, participants made 15.2 (38%) agent-dependent and 18.67 (47%) agent-independent predictions; trials that did not align with either model (6.13 trials, 15%) were included as separate regressors in the GLM.

We took a data-driven approach to identify functional regions associated with task conditions, which makes it easier to examine how multiple factors relate with signal in a given area (e.g., Satpute et al., 2013). Standardized parameter estimates of the six conditions from the first-level GLM were submitted to an omnibus ANOVA (FDR, q < .01), which resulted in 31 regions. Parameter estimates were then averaged across voxels within each region resulting in a subject-by-condition-by-regions data matrix. We then conducted a series of follow up analyses to unpack the functional profiles of these regions. First, to simplify the findings, this matrix was submitted to a k-means clustering analysis (Scikit-learn, KMeans, using the cosine distance metrics) with an elbow method cutoff calculated by computing the inertia, which is the sum of squared distances from each data point to its assigned cluster center. Data was then averaged within clusters to create a subject-by-condition data matrix per cluster.

Second, the condition-dependent response profiles were characterized by testing for differences in BOLD signal during cue v. feedback periods and agent-dependent v. agent-independent cue periods using repeated measures t-tests, and – to examine differences in feedback signals by task conditions – a 2 (prediction error v. congruent) x 2 (agent-dependent v. agent-independent prediction) repeated measures ANOVA. These analyses were conducted to characterize response profiles across regions and clusters initially identified by the omnibus F-test, which does not constrain which conditions drive variation in signal. Still, because voxel selection was based on this whole-brain test, we do not report effect size magnitudes for the fMRI analyses below, and any observed effect size indicators should be interpreted with caution due to non-independence (Vul et al., 2009).

## Results

### Behavioral Validation of the Task Paradigm

The behavioral study validated the main features of the task using a deterministic design. Figure 3 shows data from participants who completed trials with agent-dependent feedback (trials 1-20) followed immediately by the agent-independent feedback trials (trials 21-40; the opposite counterbalancing order showed parallel effects). As expected, participants first learned to make predictions consistent with the agent-dependent model (i.e., trials 1-6, Figure 3A). On trial 21, feedback switched to the agent-independent condition. Participants evidenced prediction error following the switch; performance was significantly below the 25% chance rate for the first two trials after the switch (i.e., trials 21-22), but then participants learned to make predictions consistent with agent-independent priors. Figure 3B and C show that participants also reported increased confidence and reduced surprise during the initial learning trials; the highest confidence, but also the greatest surprise at the switch trial. They then showed increasing confidence and reduced surprise with learning (trials 23+). These findings confirm that participants can flexibly adopt distinct mental models to make movement predictions.

**Figure 3.**
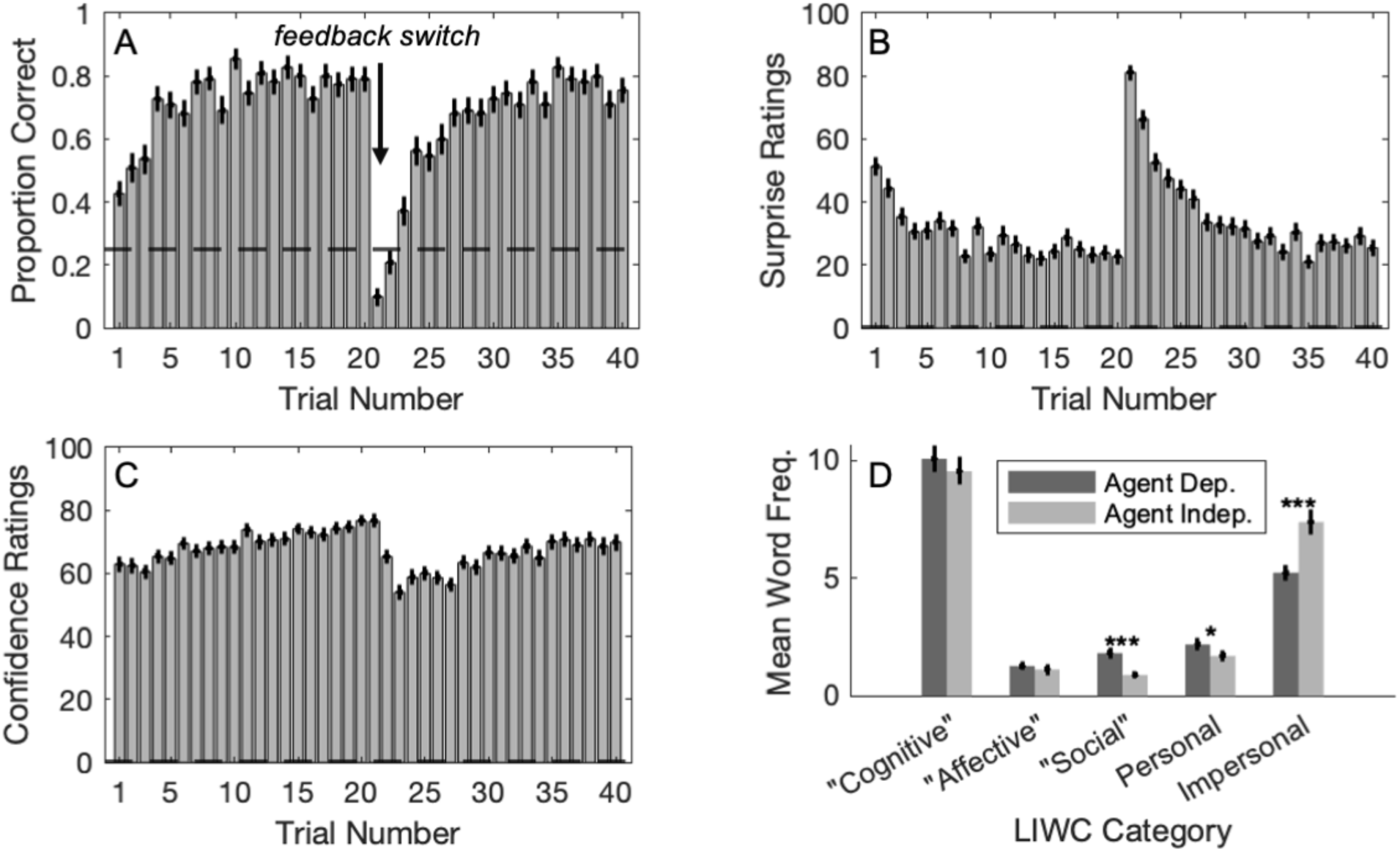
Paradigm validation. In the behavioral study, participants (N = 110) were randomly assigned to complete 20 trials of either agent-dependent or agent-independent feedback, upon which the feedback condition switched (on trial 21), for the remaining 20 trials. Results are presented averaging across counterbalancing conditions with standard error bars. (A) Participants learned to make predictions consistent with the assigned feedback condition (trials 1–4), then showed below-chance performance upon the switch to the other condition (trial 21), followed again by learning of the other condition (chance = 25%). (B) Self-reported prediction confidence [“How confident are you in your prediction?” 0 ∼ “not at all confident”, 100 ∼ “very confident”] accrued with learning, peaked on the switch trial, and then accrued with learning again. (C) Subjective surprise [“How expected was that outcome?” 0 ∼ “not at all expected”, 100 ∼ “very expected”] also tracked with learning and was greatest for feedback on the switch trial. (D) Open-ended verbal descriptions of movement included more “Social” category word use, personal pronouns, and fewer impersonal pronouns, during the block of trials with agent-dependent feedback, while “Cognitive” and “Affective” category word use was no different between conditions (LIWC text analysis; \**p <* .05, \*\*\**p* <.001).

Finally, participants provided open-ended verbal descriptions of the movement on each trial. While on average more words were used during the agent-dependent condition (t(108) = 3.70, p < 0.001), this varied significantly by semantic content. For example, while language pertaining to “cognition” and “affect” did not differ between conditions (t(108) = 1.43, p > 0.15; t(108) = 0.56, p > 0.5, respectively), participants used more “social” language in the agent-dependent condition (t(108) = 4.72, p < 0.001) as expected, as well as more personal pronouns (t(108) = 2.61, p < 0.02) and fewer impersonal pronouns (t(108) = -4.46, p < 0.001). These findings are consistent with our hypothesis that feedback would lead to the formation of two distinct mental models, and that the agent-dependent model would involve a relatively more social conceptualization.

A probabilistic task was used in the fMRI study. To confirm that participants were sensitive to using the feedback they received on prior trials to inform movement predictions on future trials, we used a sliding window analysis wherein we calculated the proportion of dominant feedback videos over a window of *k* prior trials (ranging from 1 to 10), and examined whether this predicted movement predictions on the subsequent trial using simple linear regressions. Feedback probability from past trials predicted subsequent movement predictions across several window sizes (peak at *k =* 7; *b =* 1.54, SE = .69*; p* < .05; see Figure S2).

### Brain Regions Associated with Task Parameters

We used a data-driven approach to test whether the pattern of signal across these areas reflects a common underlying activation profile during agentic movement perception, or whether it reflects functionally dissociable clusters or networks of brain regions. First, a one-way ANOVA showed that hemodynamic signals associated with task conditions were widely distributed across the cortex and cerebellum (Figure S3). As expected, this included several brain regions previously implicated in social cognition and mentalizing - including the dorsomedial and ventrolateral prefrontal cortex, lateral middle temporal gyrus, and lateral posterior parietal cortex - and also action observation, action planning, and mirroring - including the premotor cortex and lateral superior parietal lobule. While we use these psychological domain labels to refer to sets of brain regions and connect our findings to relevant prior literature in social cognitive neuroscience, we note that these regions are not necessarily exclusively involved in those domains; they may also participate in other psychological phenomena (Barrett & Satpute, 2013).

### Action Observation and Mentalizing Networks are Differentially Engaged during Prediction Formation and Feedback Processing

Next, a k-means clustering analysis revealed three clusters of brain regions with different response profiles across conditions. Notably, clusters dissociated along the lines of previously observed mirroring and mentalizing networks for clusters 1 and 2 (Figure 4, top), respectively, while cluster 3 was composed of other posterior cortical areas (Figure S4, top). The data-driven approach also revealed functionally heterogeneous contributions of cerebellum; each cluster involved different sectors of cerebellum along with cortical areas. We focused our analyses on clusters 1 and 2 given their established role in social cognition (descriptive information on cluster 3 is provided for completeness).

**Figure 4.**
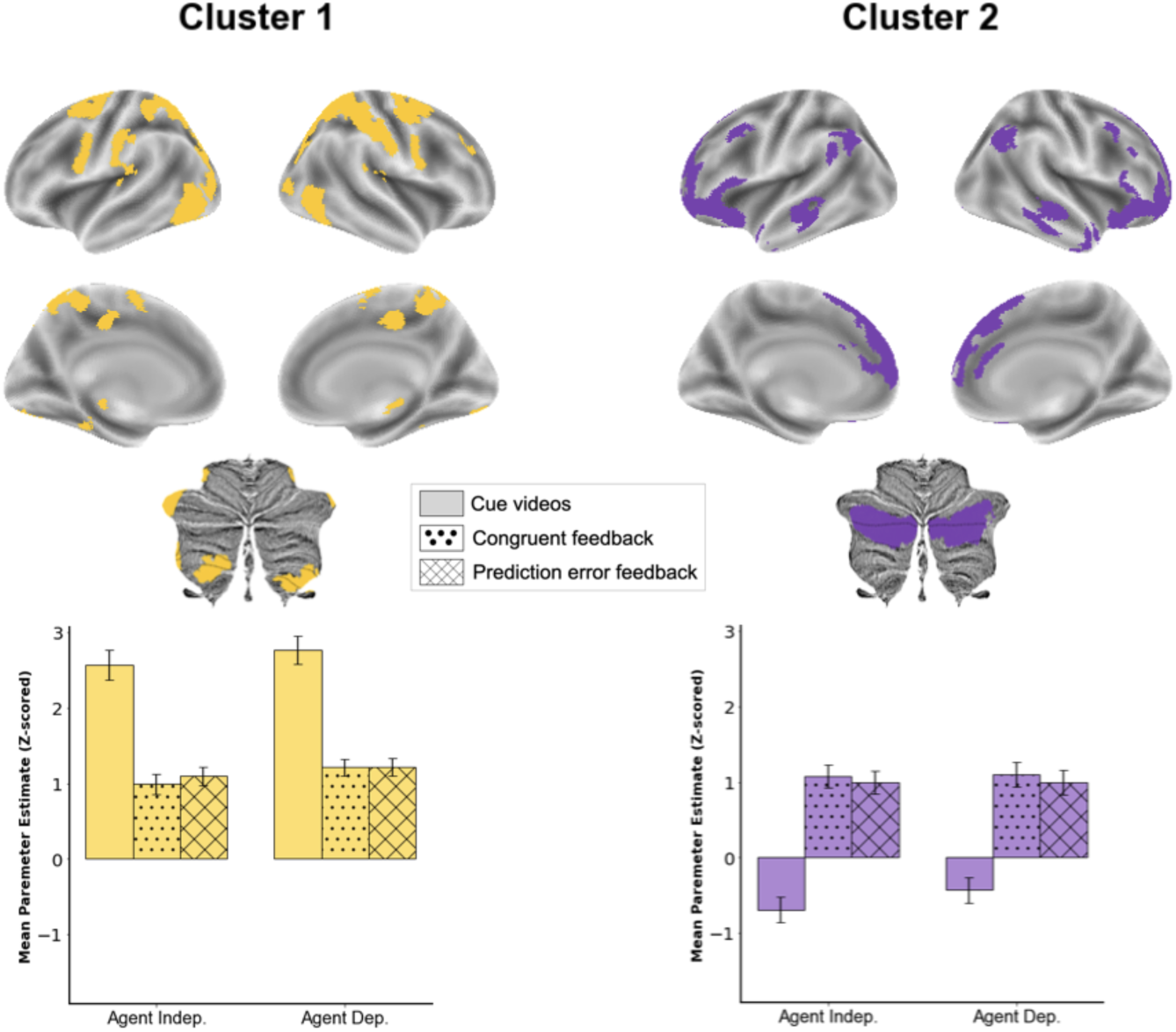
Brain regions clustered based on similarity of response profiles across task conditions. Cluster 1 consisted of areas previously associated with action observation and had greater signal during the cue periods than feedback periods. Cluster 2 consisted of areas previously associated with theory of mind and mentalizing had signal below baseline during the cue period and signal above baseline during the feedback period. Cluster 3 (not shown here, but see Figure S4) generally had reduced signal across conditions and included the posterior mid-cingulate cortex and portions of the occipital cortex. Cerebellum is displayed using flat maps (Diedrichsen & Zotow, 2015). Cluster 1 included hemispheric lobules VIIIA, VIIIB, and IX; cluster 2 included crus I and II.

As shown in Figure 4 (bottom), the functional activation profile of Cluster 1 (action observation and mirroring) had significantly greater signal during the cue period than the feedback period (*t*(29) = 14.221, *p* < 4.7 × 10⁻¹⁴), whereas cluster 2 (theory of mind and mentalizing) showed the opposite pattern with greater signal during the feedback period and reduced signal during the cue period (*t*(29) = -11.375, *p* < 1.2 × 10⁻¹¹). Notably, although these findings support a functional dissociation between the clusters, the voxel selection was based on a whole-brain analysis. As a result, we do not report effect size magnitudes for this or other fMRI analyses presented below, and any observed effect size indicators should be interpreted with caution due to non-independence (Vul et al., 2009).

Next, we compared signal during the cue-period for agent-dependent v. agent-independent trials to see if functional activity differed based on the type of predictions participants were making (i.e., if they made agent-dependent or agent-independent predictions). There were no significant differences in any of the clusters (|*t| <* 1.3, *p*s > .15), suggesting that functional activity in this task during the cue period generalized across agent-dependent and agent-independent conditions. In the cerebellum, cluster 1 included vermian regions (HVIIIa and HVIIIb), while cluster 2 included crus I and II. Notably, these areas have been functionally connected to regions implicated in action observation, and in mentalizing/theory of mind, respectively, in previous studies examining resting state functional connectivity (Diedrichsen & Zotow, 2015).

Because the main analysis focussed on differences between task conditions that all involved video watching, we conducted an additional analysis to identify regions engaged in visual processing regardless of task conditions. Most visually responsive regions were not modulated by task manipulations - suggesting that several brain regions were sensitive to visual processing irrespective of any differences in the visual stimuli related to task conditions (Figure S7). A subset of areas—primarily in cluster 1, including the parietal cortex, lateral occipital cortex, and premotor cortex— while showing greater activity during visual processing in general also showed greater activation during cue than feedback periods. The selective modulation of these areas suggests these regions are not merely responsive to visual input, but are additionally engaged during the formation of predictions about upcoming movements, consistent with their involvement in action observation and preparatory processes.

### Condition-Dependent Prediction Error Signals

Finally, we examined signal associated with processing congruent v. incongruent feedback. Here, we tested whether modulation by feedback type was content general or content dependent using 2 (prediction content: agent-dependent or agent-independent) x 2 (feedback type: congruent or incongruent) repeated measures ANOVAs. Interaction effects tend to be more subtle than main effects and hence may cancel each other out when examined at the cluster level; hence, analyses were conducted at the region level. Of the brain regions showing modulation during feedback (presented in Figure 5), all showed an interaction effect wherein congruent and incongruent feedback depended on condition. For some of these areas, the difference between incongruent and congruent feedback appeared to be greater when participants made agent-dependent predictions, including activation spanning the nucleus accumbens and subgenual anterior cingulate cortex, bilateral somatosensory cortex, and the right dorsolateral prefrontal cortex regions. Other areas showed this difference to be greater when participants made agent-independent predictions, including in bilateral parahippocampal and fusiform cortex. Details for each region are included in Figure S5 and Table S1. Overall, these findings suggest that activity related to prediction error was dominated by content-dependent signals.

**Figure 5.**
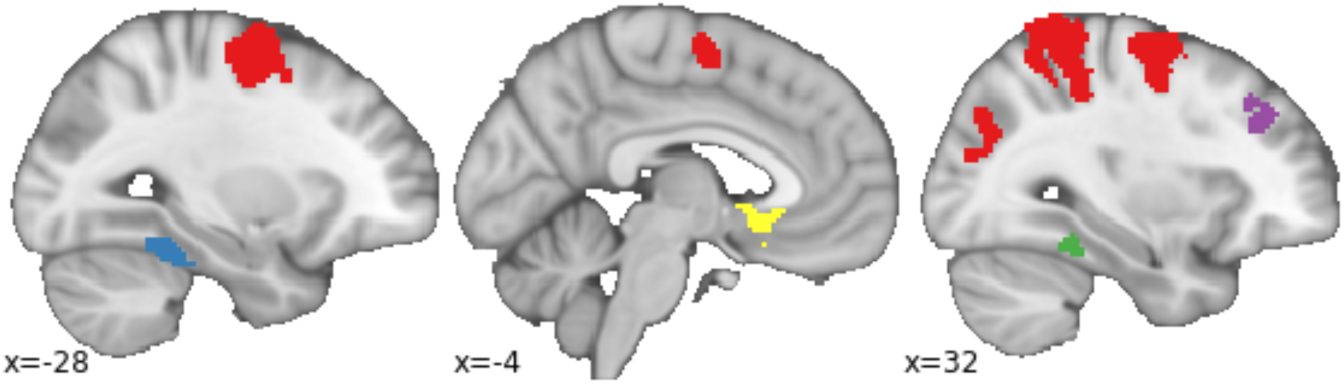
Functional activity during feedback type (incongruent v. congruent) depended on the type of mental model. Some regions—such as the nucleus accumbens and subgenual anterior cingulate (yellow), bilateral somatosensory cortex (red), and right dorsolateral prefrontal cortex (purple)—showed stronger incongruence effects when predictions consistent with agent-dependent priors were made, while others—such as the bilateral parahippocampal cortex and fusiform gyrus (blue and green) —showed stronger incongruence effects when predictions consistent with agent-independent priors were made.

## Discussion

In this study, we observed a functional dissociation across neural networks involved in social cognition. Brain regions previously implicated in action observation, motor planning, and mirroring - including the lateral parietal cortex and premotor cortex - showed heightened activity during cue processing, when participants formulated specific predictions about agentic movement. In contrast, regions previously implicated in theory of mind and mentalizing - including the dorsomedial and ventral lateral prefrontal cortex, temporoparietal area, and lateral temporal cortex - all showed greater activity during feedback processing, when participants could use feedback to update relatively abstract, agent-dependent and agent-independent priors. A parallel functional dissociation was observed in the cerebellum with different portions joining these respective networks. Additionally, we found that incongruent feedback, or prediction error, was represented in a condition-dependent way; that is, different areas were associated with incongruent feedback depending on whether an agent-dependent vs. agent-independent prediction was made. These findings highlight how distinct brain systems contribute to different phases of social prediction, offering new insights into the mechanistic functional roles of these networks in supporting social cognition.

### Predictive Processing Theories and Social Cognitive Neuroscience

While numerous studies have identified where social processing occurs in the brain, the cognitive mechanisms supported by these regions—particularly from a neurally plausible, computational perspective—remain unclear (Adolphs, 2009).

Predictive processing offers a promising framework to address this gap (Kilner et al., 2007), positing that the brain maintains a generative model of the sensory environment and uses deviations from predicted input—prediction errors—to update that model (Clark, 2013; Friston, 2005). According to hierarchical predictive processing models, lower-level sensory areas generate fine-grained, modality-specific predictions, while higher-level association areas generate more abstract, content-dependent predictions (Adams et al., 2015; Bastos et al., 2012; Rao & Ballard, 1999). In our task, participants predicted agentic movement by integrating visual information from cue videos with higher-level mental models—specifically, agent-dependent and agent-independent interpretations (Schultz et al., 2005). We observed a functional dissociation across brain regions, with areas associated with action observation and mirroring showing greater activation during cue processing, and regions implicated in theory of mind and mentalizing showing greater activation during feedback. This dissociation may reflect distinct roles in prediction and model updating: cue-related activity may reflect the generation of a prior or mental model to guide prediction, whereas feedback-related activity may reflect the incorporation of prediction error to update that model.

Areas previously implicated in action observation and action planning—including the intraparietal sulcus, premotor cortex, and lateral occipital cortex—are hierarchically situated below areas associated with mentalizing (Margulies et al., 2016; Paquola et al., 2025), and may issue predictions about expected movements based on immediate visual cues and contextual information (Cerliani et al., 2022; Kilner et al., 2007). In contrast, areas previously implicated in mentalizing—including the dorsomedial and ventrolateral prefrontal cortex, temporoparietal junction, and lateral temporal cortex—may contribute to updating these predictions based on more abstract, conceptual information (Koster-Hale & Saxe, 2013; Satpute & Lindquist, 2019; Smallwood et al., 2021; Spunt et al., 2016). Feedback signals modulated by prediction error are expected to recruit these higher-order areas to revise the generative model. Together, these findings are consistent with a distributed and hierarchical account of predictive processing in social cognition (Brown & Brüne, 2012; Heil et al., 2019; Köster et al., 2020). Concrete, contextualized predictions are driven by lower-level sensorimotor systems, while abstract model revision is supported by systems that monitor relational structures and track feedback over time.

### The Cerebellum and Social Cognition

A growing body of research implicates the cerebellum in social cognition. Neuroimaging studies have shown that cerebellar regions, particularly in the posterior lobules, are more active during social compared to non-social tasks (Guell et al., 2018; Jack & Pelphrey, 2015; Van Overwalle et al., 2014); c.f. (Metoki et al., 2022), and neuropsychological findings link cerebellar damage to deficits in social reasoning and mental state attribution (Clausi et al., 2019; Tamaš et al., 2021), which may vary with development (Olson et al., 2023). While neuroimaging and neuropsychological data suggest cerebellar involvement in social cognition generally, a more specific hypothesis prominently featured in a recent consensus paper posits that the cerebellum “facilitates social cognition by supporting optimal predictions about imminent or future social interaction and cooperation” (Van Overwalle et al., 2020, p. 833). Support for this predictive role comes primarily from computational and neuroanatomical models (e.g., Doya, 1999; Sokolov et al., 2017; Wolpert et al., 2003) and from studies in non-social domains, such as motor control and sequence learning.

Our study builds on this theoretical foundation by directly testing the cerebellum’s role in prediction within a social cognitive task, bridging a key empirical gap in the literature. Specifically, our task design isolates two components: the cue period, associated with deploying predictive mental models, and the feedback period, linked to updating these models based on new information. Previous studies using animated shape videos were not designed to disentangle these processes. By isolating these phases of our task, we found that distinct cerebellar regions were selectively engaged during each stage. Consistent with prior work, crus I and II—previously implicated in social cognition and identified as part of the default mode network (DMN) through resting-state fMRI (Buckner et al., 2011)—were robustly engaged during our task.

Notably, these regions were clustered with canonical DMN and mentalizing areas (cluster 2), based on their shared task-evoked profile. Unlike earlier studies that broadly linked these areas to social processing, we found that they were less active during cue periods (below fixation baseline) and more active during feedback periods, suggesting a specialized role in updating mental models based on abstract priors (e.g., agent-dependent vs. agent-independent). This functional dissociation points to a role for crus I/II and their associated cortical partners in revising social predictions rather than generating them.

In contrast, lobules VIIIA, VIIIB, and IX showed greater activity during cue periods than during feedback, suggesting a distinct functional role from Crus I/II. These regions are affiliated with different large-scale networks: VIIIA is part of the somatomotor network, VIIIB aligns with the ventral attention network, while both of those networks overlap with the action observation network. In our study, they clustered together in cluster 3, which included cortical regions previously implicated in action observation and mirroring processes. The increased activity in these lobules during cue processing suggests a role in generating sensory-driven predictions based on local visual cues, potentially supporting lower-level, embodied aspects of social perception. This distinction aligns with a broader framework in which the cerebellum contributes to social cognition through multiple predictive mechanisms—ranging from abstract mental model updating to more concrete predictions of expected sensory information – with levels of abstraction in prediction potentially determined by parallel cortical-cerebellar networks.

### Limitations and Future Directions

The present study is limited in several ways that may be addressed in future work. First, the novel fMRI task offers unique opportunities for computational modeling of behavior. For example, such models could be used to estimate the strength or confidence of agent-dependent and agent-independent priors. We chose not to apply these techniques here due to the limited number of trials, but future work with more extensive trial counts may allow for more robust modeling approaches.

Second, the task used a probabilistic design to influence the strength of a prior. While this design was effective in shaping participants’ behavioral predictions, it’s possible that participants continued to represent both priors to some degree, which may have reduced our ability to detect differences in neural activity during the cue period that are specific to agent-dependent vs. agent-independent models. Future work using a reversal learning design, in which the strength of one prior increases while the other decreases across blocks, may help clarify how these models are differentially acquired and represented in the brain.

Third, we examined agentic movement using an animated shapes task. While these stimuli are less naturalistic than images or videos involving real people, prior work using the classic animated shapes videos or variants thereof (Castelli et al., 2004; Gobbini et al., 2007; Martin & Weisberg, 2003; Wheatley et al., 2007) has shown that people spontaneously make social inferences from these stimuli, and also that they reliably engage many of the same brain regions implicated in mentalizing and theory of mind as when using externally valid stimuli (Jacoby et al., 2016; Schurz et al., 2014; Spunt et al., 2011). For our purposes, the use of abstract stimuli was particularly useful because it allowed us to examine the flexible deployment of priors in making predictions about agentic movement—which can be obscured when using more naturalistic stimuli that come imbued with semantic meaning. While this approach offers advantages in terms of internal validity for isolating specific mechanisms, and the neural correlates we observed were consistent with studies using more naturalistic paradigms, future work should nonetheless explore the generalizability of these findings to everyday social contexts (for discussions on predictive processing theory and external validity, see Lee et al., 2021; Miller et al., 2019).

## Summary and Conclusions

Much research in social neuroscience has focused on which brain regions are engaged when processing social content, yet it remains unclear what these areas are doing in terms of underlying mechanisms. Here, we approached this question using predictive processing theory, which suggests that the brain instantiates a generative model of its sensory environment. We developed and deployed a novel fMRI task that separates into stages the processes involved in forming a mental model for making prediction of agentic movement from those involved in processing feedback. Brain regions previously associated with action observation/mirroring along with cerebellar lobules VIIIA, VIIIB, and IX, were relatively more engaged when processing cue videos to make predictions v. processing feedback videos to update predictions on future trials, whereas areas previously implicated in mentalizing/theory of mind along with cerebellar crus I and II showed the opposite pattern. Further, we found that greater prediction errors (i.e., incongruent v. congruent feedback) was represented in a condition-dependent manner. These findings offer new insights into how action observation/mirroring and mentalizing networks - involving both cortical and cerebellar areas - play complementary roles in supporting social cognition, more broadly, from a predictive processing perspective. Future work may use this task paradigm to further investigate the computational and mechanistic properties of these networks and how they vary in relation to individual differences in neurotypical and clinical populations, and across the lifespan.

## Acknowledgements

This research was supported by the Brain and Cognitive Sciences Division of the National Science Foundation [award number: 2241938]. We thank Stephanie Fiedler, Ellie Diederich, Maya Sundel, Haden Pelletier, and Eric Feng for assistance with stimuli creation and data collection including

## Data and Code Availability

Anonymized data will be deposited in OpenNeuro (https://openneuro.org/) upon publication. The stimulus set is available in Github at https://github.com/yiyuwang/SocialPrediction_AnimatedShapesTest

Analysis scripts are available in Github at https://github.com/yiyuwang/SocialPrediction

## Supplementary

**Figure S1.**
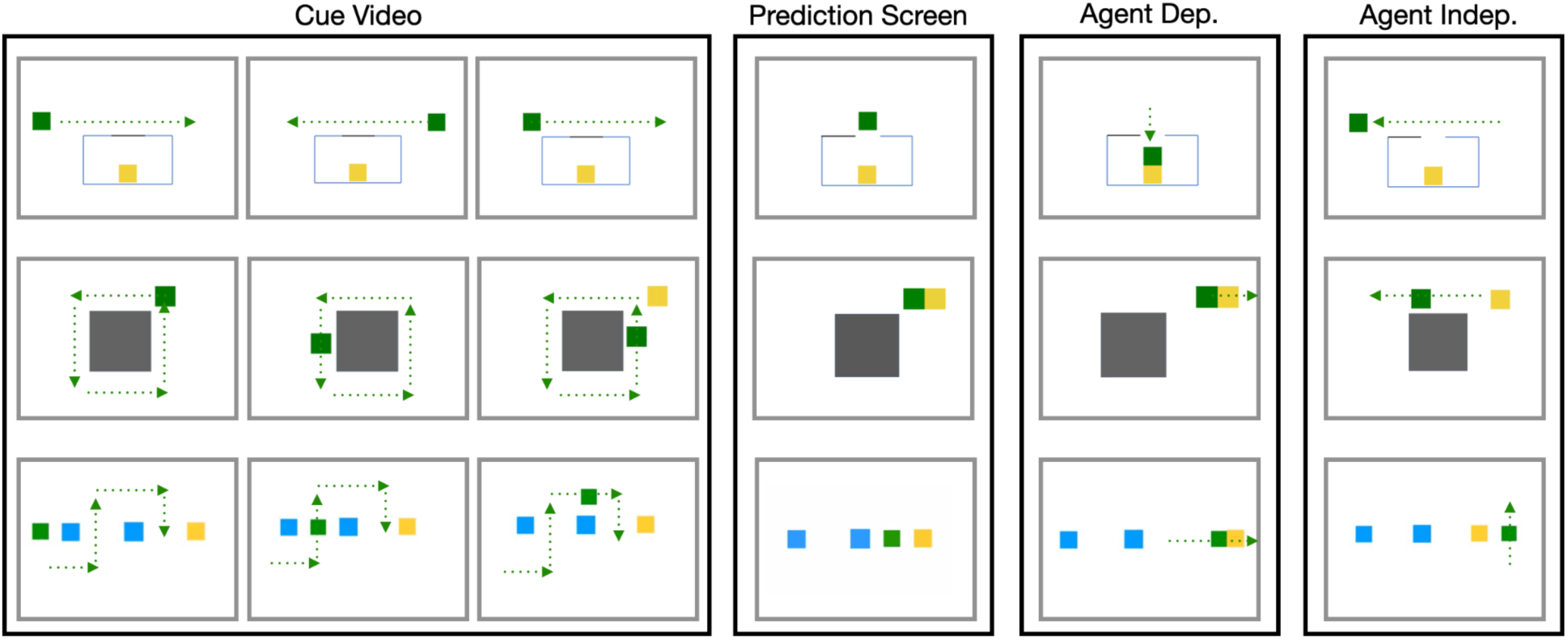
sample videos

**Figure S2.**
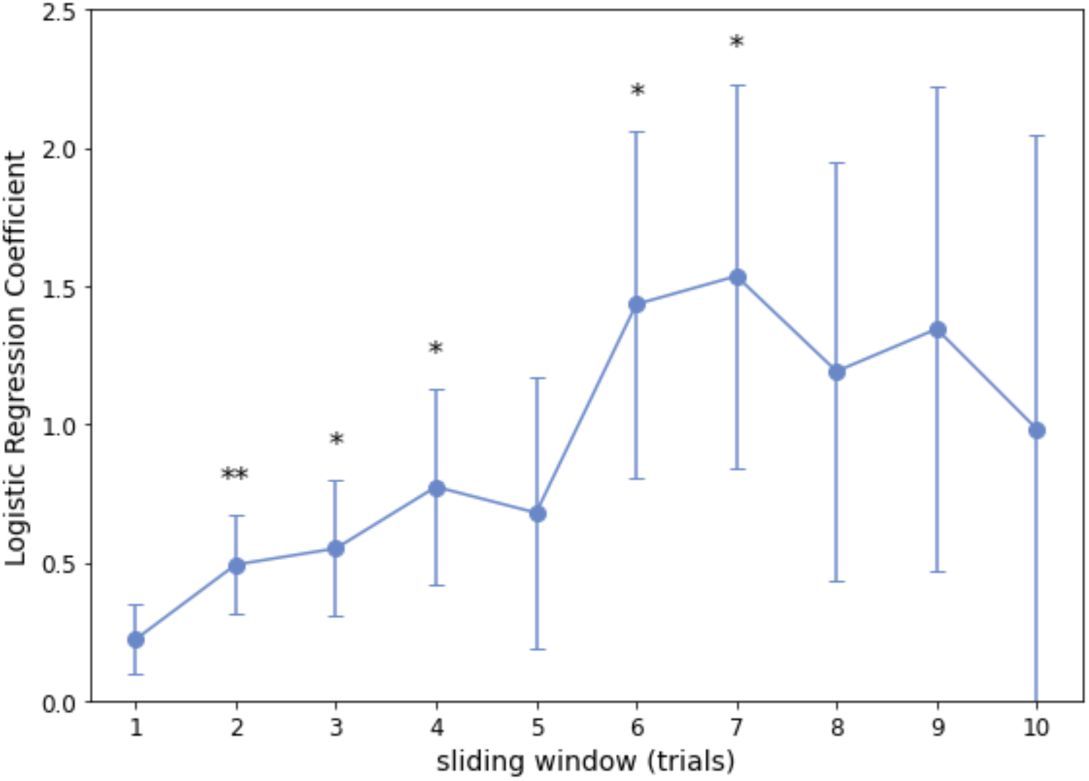
Proportion of feedback type influences predictions on subsequent trials. Participants completed a probabilistic task in the fMRI study. A sliding window analysis was conducted to test whether participants were sensitive to using the feedback on prior trials to inform behavioral predictions on future trials. We calculated the proportion of dominant feedback videos over a window of k prior trials (ranging from 1 to 10, x-axis), and examined whether this predicted predictions on the subsequent trial using simple linear regressions. Feedback probability from past trials predicted subsequent movement predictions across several window sizes. Error bars reflect standard errors of the mean. * p < 0.05, ** p < 0.01

**Figure S3.**
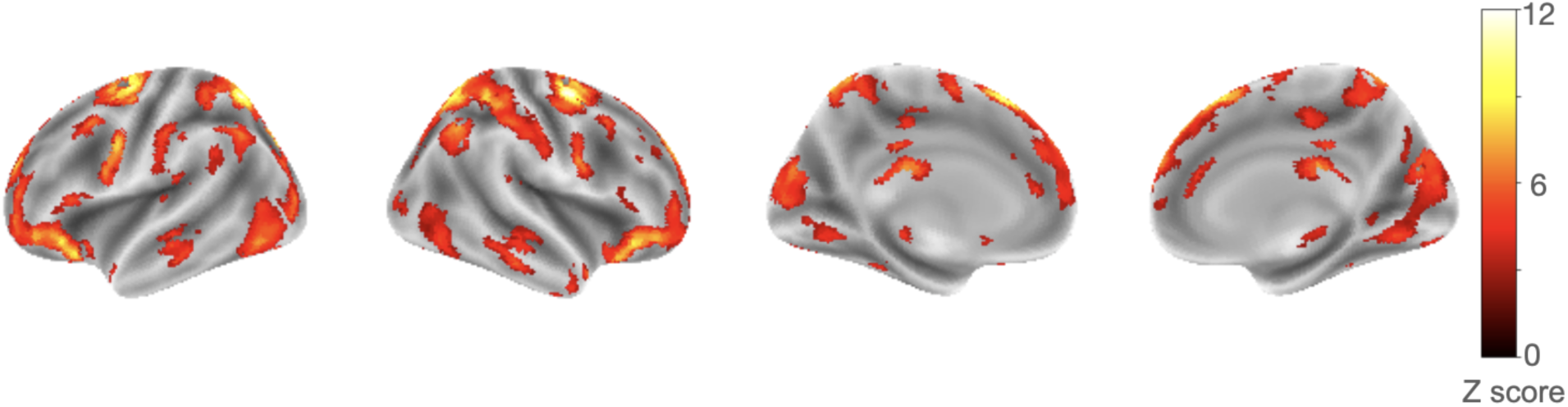
Brain regions associated with task variables. A one-way ANOVA revealed brain regions with functional activity associated with task conditions (FDR corrected at q<0.05). Parameter estimates across voxels averaged across voxels, per region, per subject. These values were then submitted to a clustering analysis to organize regions into clusters of activation profiles for analysis (Figure 4 and Figure S4).

**Figure S4.**
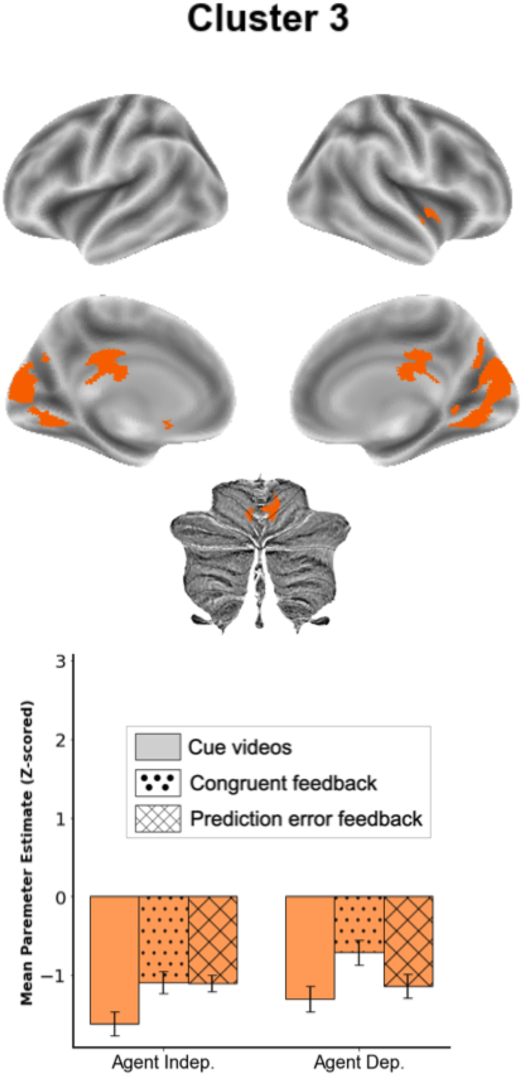
Unlike clusters 1 and 2, cluster 3 - which included the posterior mid-cingulate cortex and portions of the occipital cortex - had reduced signal across conditions.

**Figure S5.**
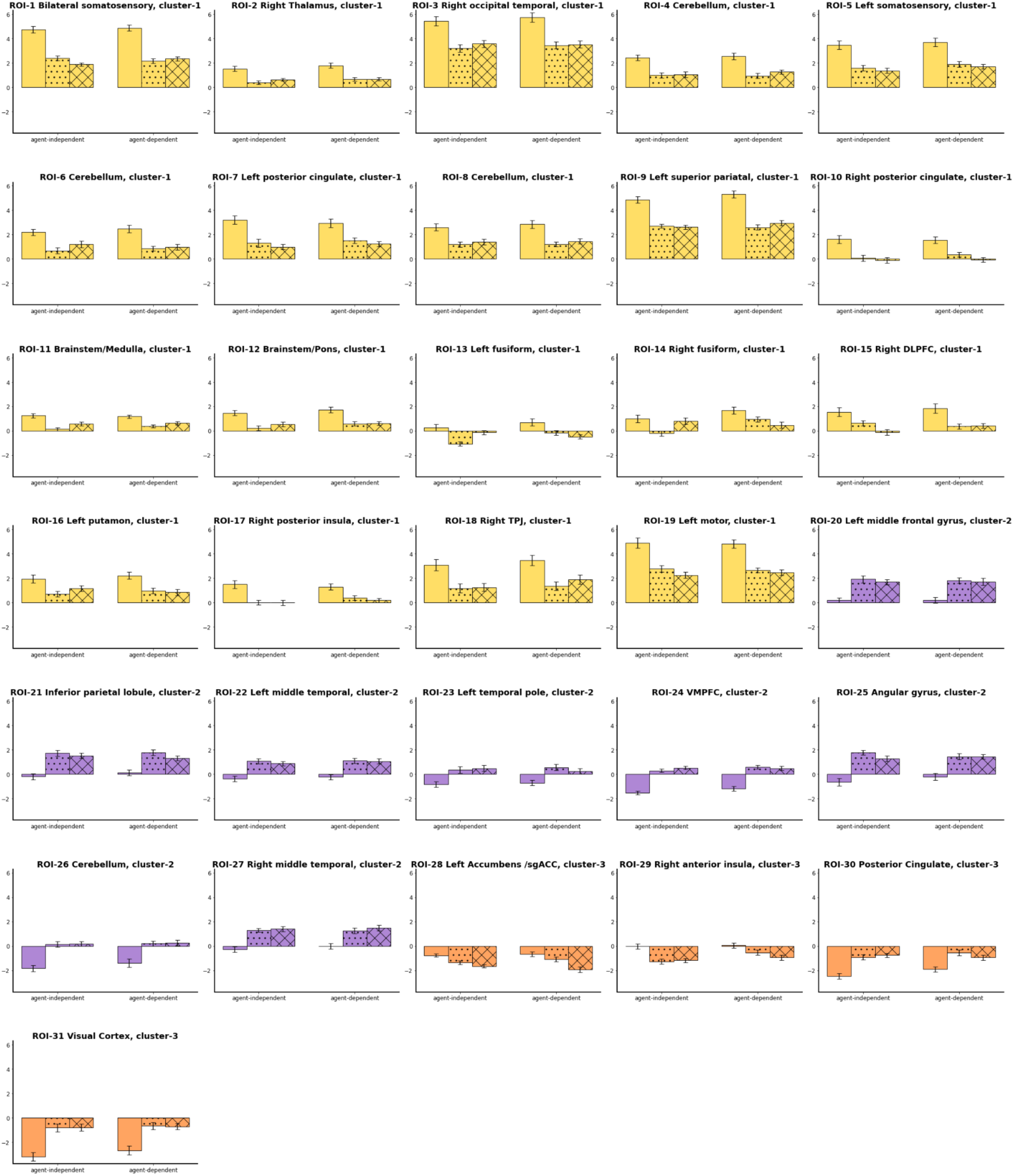
Activation profiles for each region, sorted and color-coded according to cluster (cluster 1 ∼ yellow; cluster 2 ∼ purple; cluster 3 ∼ orange). The activation profile for clusters 1 and 2 (as shown in Figure 4) provided a reasonable summary of the individual regions assigned to those clusters. There was greater heterogeneity for cluster 3; while all regions showed at or below baseline signal across conditions, for some regions signal was lower during the cue period while for others it was lower during the feedback period.

**Figure S6.**
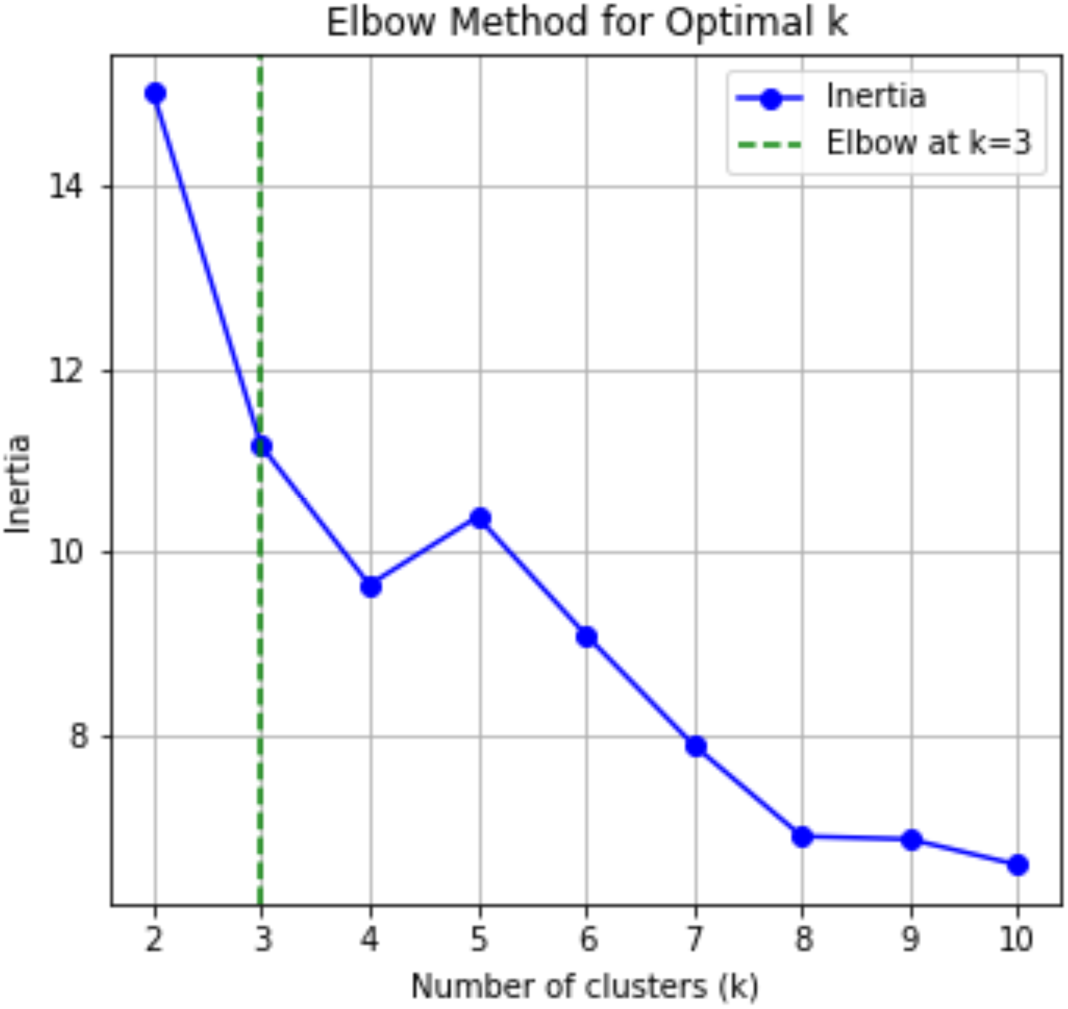
An optimal *k* value was determined at k = 3 using the elbow method based on inertia scores for different clustering solutions. Of note, inspection of the activation profiles in Figure S5 suggests that, at least for clusters 1 and 2, the cluster-level summary provides a reasonable proxy for the response profiles across individual regions.

**Figure S7.**
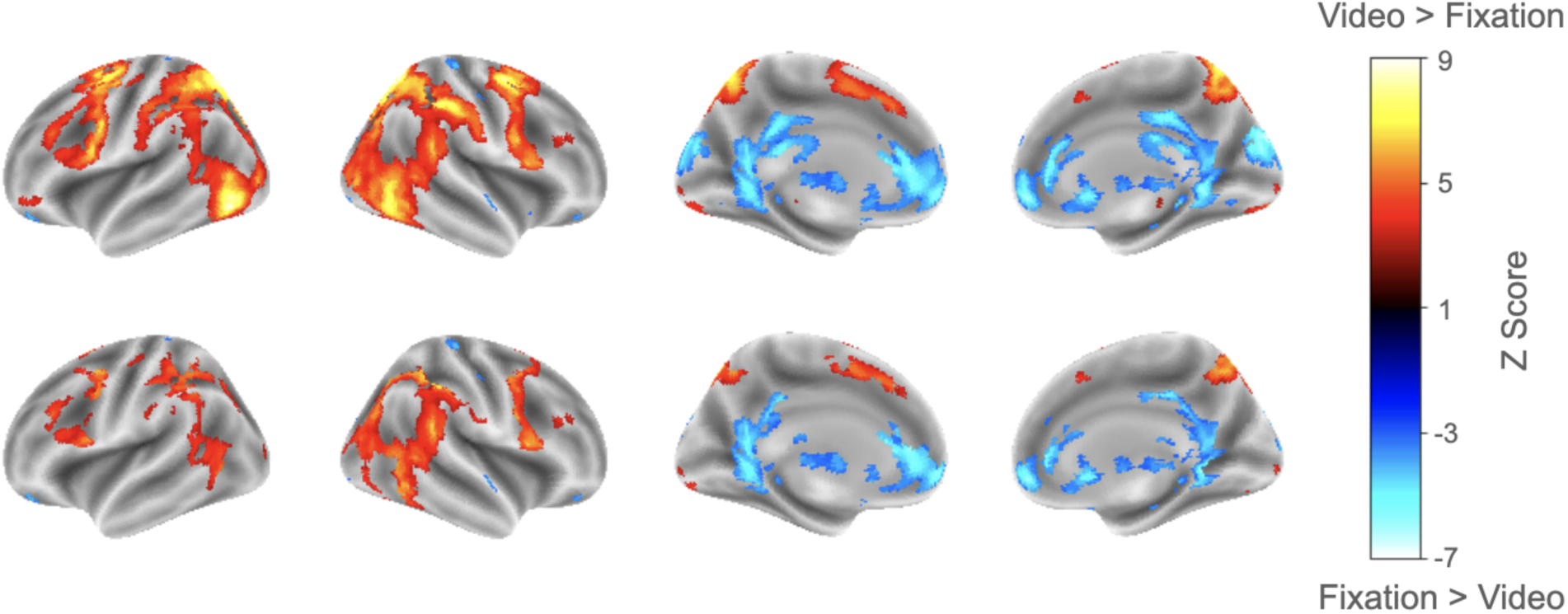
Activity associated with video watching. *Top row*: Functional activity during video periods relative to fixation baseline was observed in the posterior and lateral occipital cortex, parietal lobule, and frontal cortex. Decreased activity was noted in the ventromedial prefrontal cortex, posterior cingulate cortex, retrosplenial cortex, and subcortical regions. *Bottom row*: To isolate regions engaged in visual processing but not modulated by cue vs. feedback baseline or other task conditions, we masked the above contrast using the omnibus F-test from the main manuscript. Most visually responsive regions identified by the [video – fixation baseline] contrast were not sensitive to the specific task manipulations. However, some areas —primarily overlapping with cluster 1 (including in the parietal cortex, lateral occipital cortex, and premotor cortex) —also showed cue > feedback effects. This pattern suggests these regions contribute not only to visual processing, but also to processes specifically engaged during cue periods that may support generating predictions about upcoming movement.

**Table S1.**
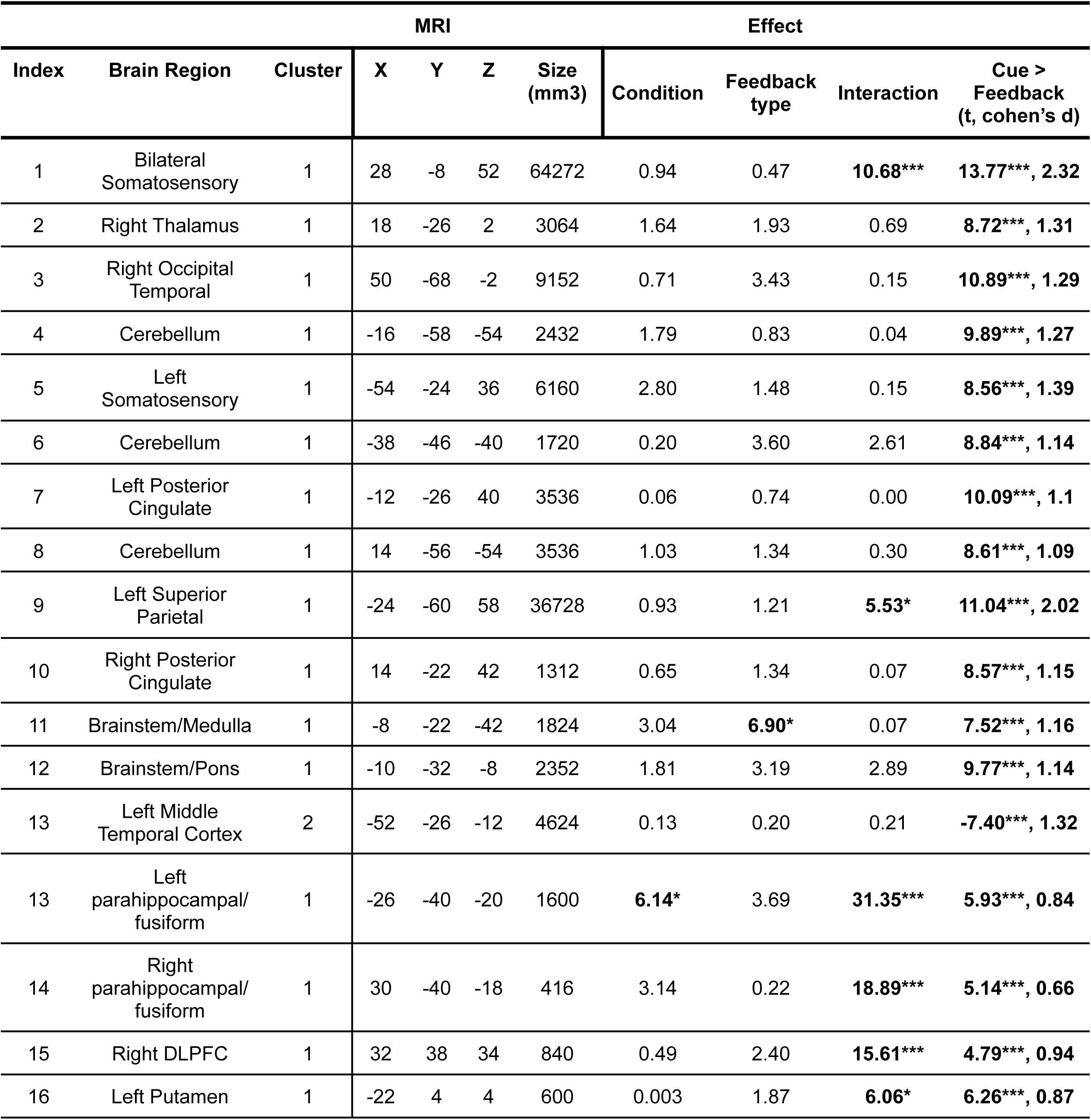
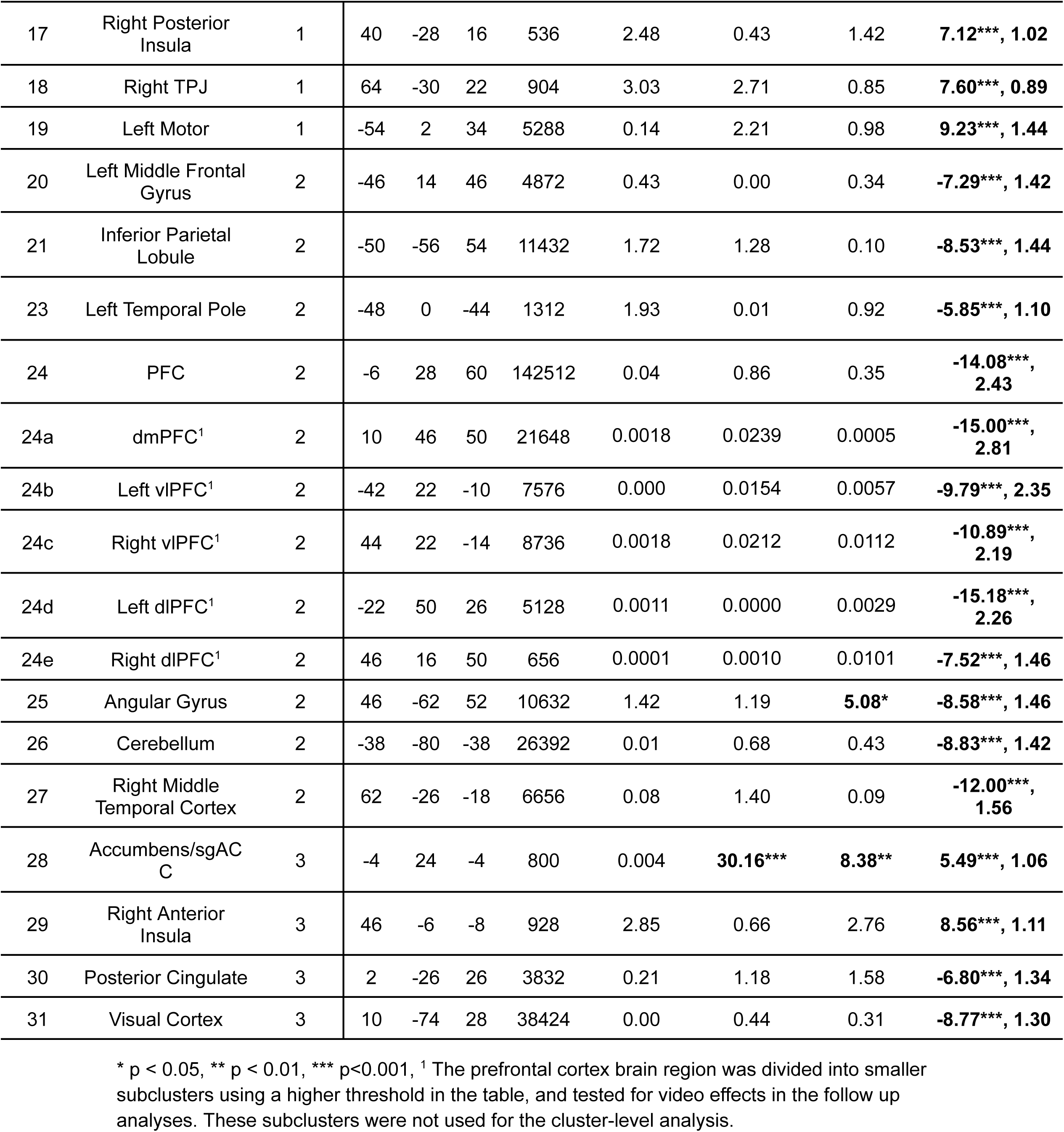
Full table of all ROIs and reported analysis effects.

