## Supplemental for "Disentangling Prediction and Feedback in Social Brain Networks: A Predictive Processing Approach"

### Supplementary materials

Figure S1 - sample videos

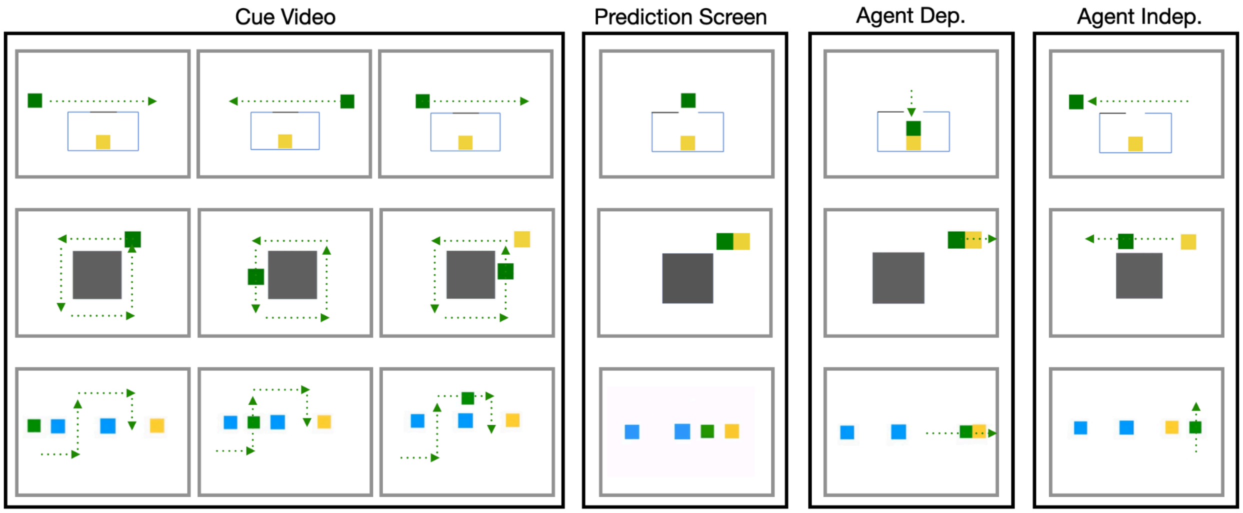

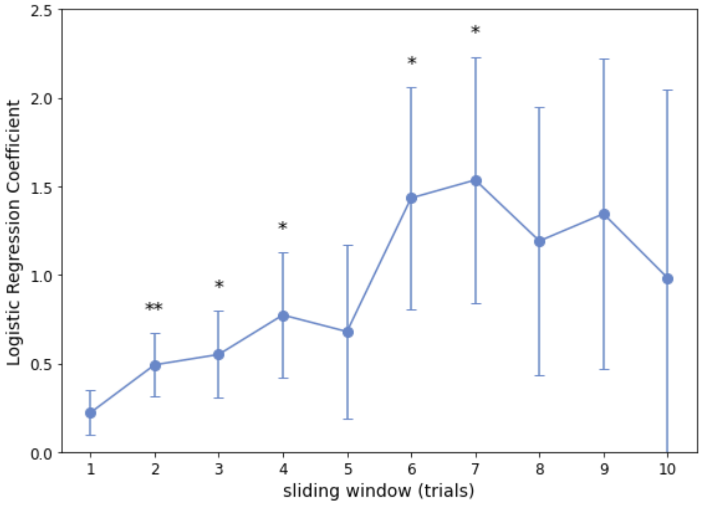


Figure S2. Proportion of feedback type influences predictions on subsequent trials. Participants completed a probabilistic task in the fMRI study. A sliding window analysis was conducted to test whether participants were sensitive to using the feedback on prior trials to inform behavioral predictions on future trials. We calculated the proportion of dominant feedback videos over a window of k prior trials (ranging from 1 to 10, x-axis), and examined whether this predicted predictions on the subsequent trial using simple linear regressions. Feedback probability from past trials predicted subsequent movement predictions across several window sizes. Error bars reflect standard errors of the mean. * $p < 0.05$, ** $p < 0.01$

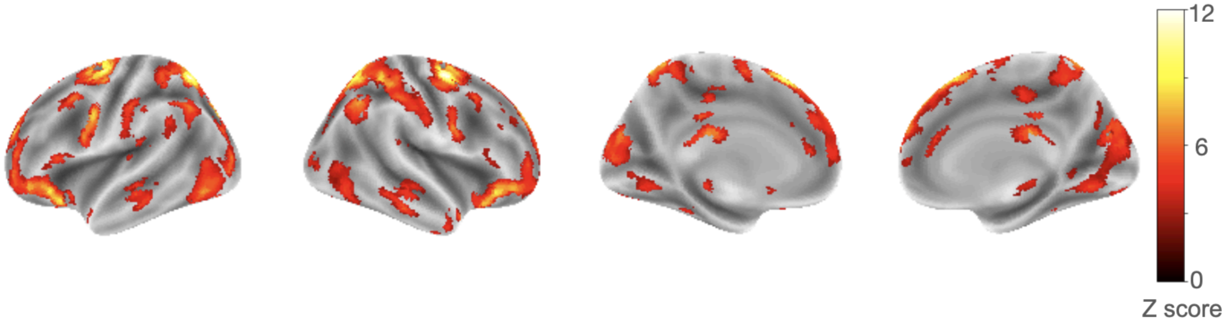


Figure S3. Brain regions associated with task variables. A one-way ANOVA revealed brain regions with functional activity associated with task conditions (FDR corrected at q<0.05). Parameter estimates across voxels averaged across voxels, per region, per subject. These values were then submitted to a clustering analysis to organize regions into clusters of activation profiles for analysis (Figure 4 and Figure S4).

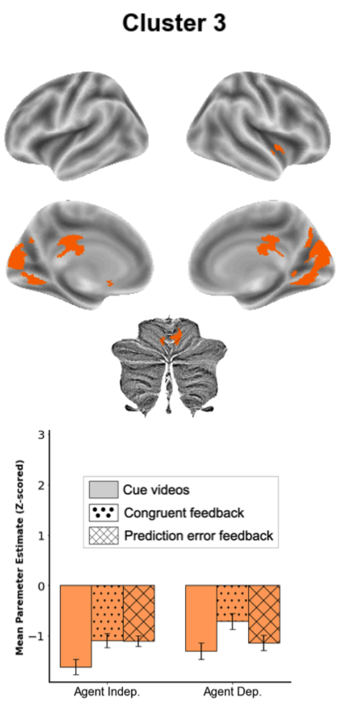


Figure S4. Unlike clusters 1 and 2, cluster 3 - which included the posterior mid-cingulate cortex and portions of the occipital cortex - had reduced signal across conditions.

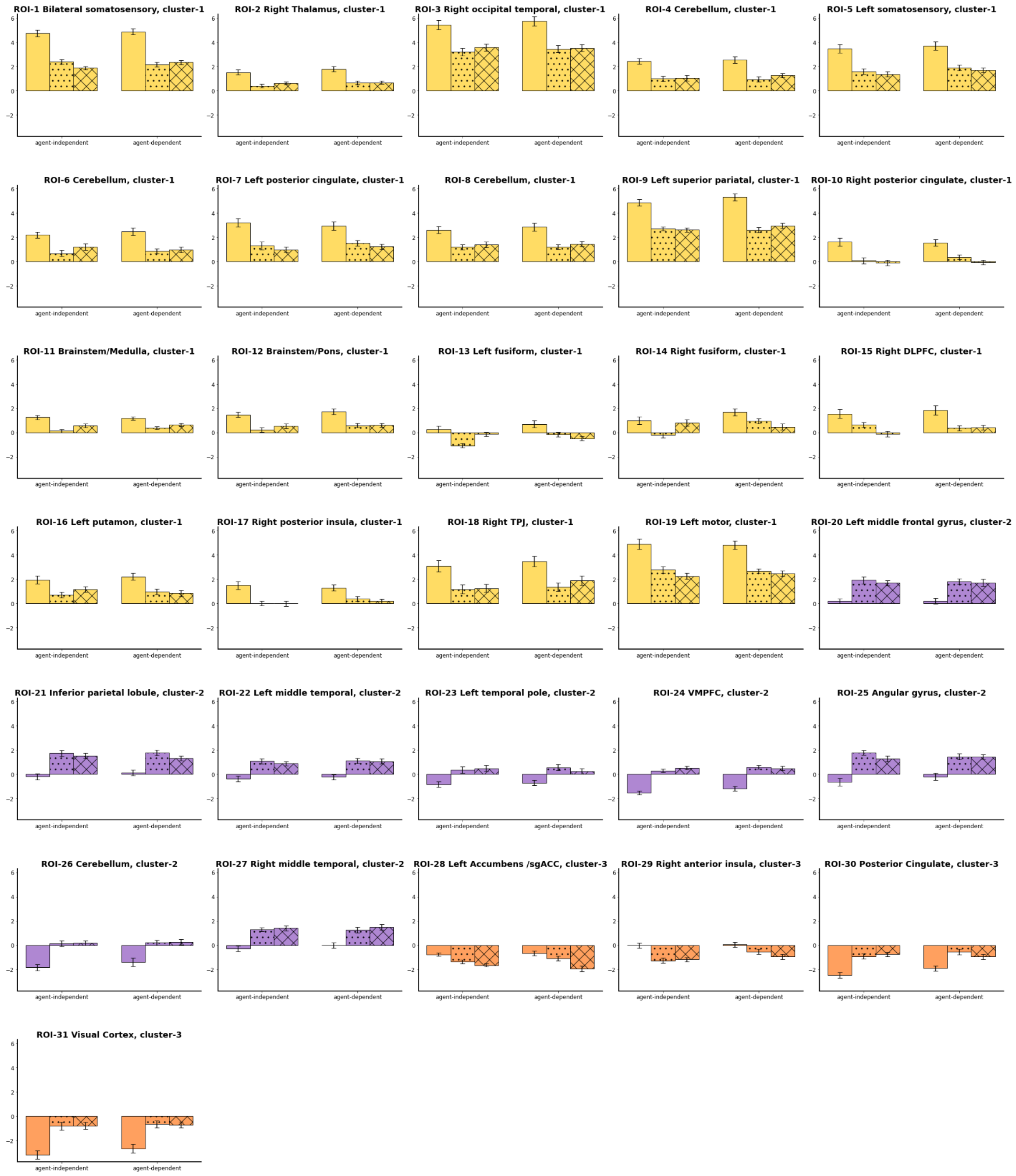


Figure S5. Activation profiles for each region, sorted and color-coded according to cluster (cluster 1 ~ yellow; cluster 2 ~ purple; cluster 3 ~ orange). The activation profile for clusters 1 and 2 (as shown in Figure 4) provided a reasonable summary of the individual regions assigned to those clusters. There was greater heterogeneity for cluster 3; while all regions showed at or below baseline signal across conditions, for some regions signal was lower during the cue period while for others it was lower during the feedback period.

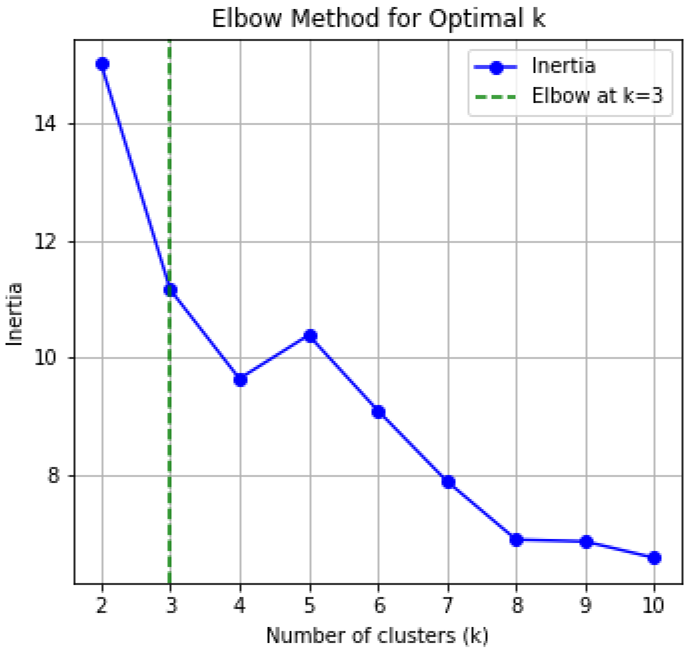


Figure S6. An optimal *k* value was determined at k = 3 using the elbow method based on inertia scores for different clustering solutions. Of note, inspection of the activation profiles in Figure S5 suggests that, at least for clusters 1 and 2, the cluster-level summary provides a reasonable proxy for the response profiles across individual regions.

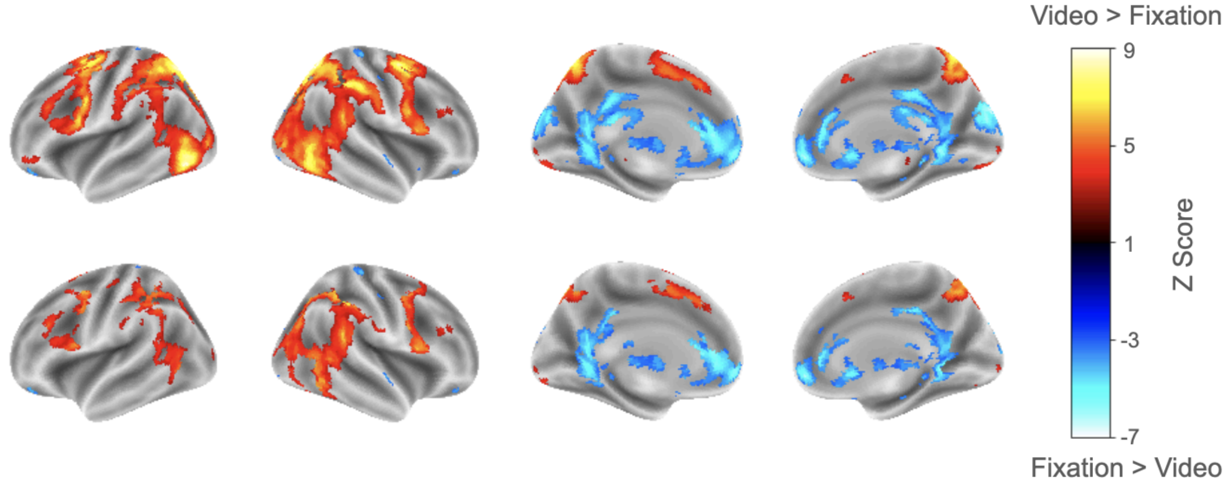


Figure S7. Activity associated with video watching. *Top row*: Functional activity during video periods relative to fixation baseline was observed in the posterior and lateral occipital cortex, parietal lobule, and frontal cortex. Decreased activity was noted in the ventromedial prefrontal cortex, posterior cingulate cortex, retrosplenial cortex, and subcortical regions. *Bottom row*: To isolate regions engaged in visual processing but not modulated by cue vs. feedback baseline or other task conditions, we masked the above contrast using the omnibus F-test from the main manuscript. Most visually responsive regions identified by the [video – fixation baseline] contrast were not sensitive to the specific task manipulations. However, some areas —primarily overlapping with cluster 1 (including in the parietal cortex, lateral occipital cortex, and premotor cortex) —also showed cue > feedback effects. This pattern suggests these regions contribute not only to visual processing, but also to processes specifically engaged during cue periods that may support generating predictions about upcoming movement.

Table S1. Full table of all ROIs and reported analysis effects.

| | | | MRI | | | | Effect | | | |
|---|---|---|---|---|---|---|---|---|---|---|
| Index | Brain Region | Cluster | X | Y | Z | Size (mm3) | Condition | Feedback type | Interaction | Cue > Feedback (t, cohen's d) |
| 1 | Bilateral Somatosensory | 1 | 28 | -8 | 52 | 64272 | 0.94 | 0.47 | **10.68*** ** | **13.77***, 2.32** |
| 2 | Right Thalamus | 1 | 18 | -26 | 2 | 3064 | 1.64 | 1.93 | 0.69 | **8.72***, 1.31** |
| 3 | Right Occipital Temporal | 1 | 50 | -68 | -2 | 9152 | 0.71 | 3.43 | 0.15 | **10.89***, 1.29** |
| 4 | Cerebellum | 1 | -16 | -58 | -54 | 2432 | 1.79 | 0.83 | 0.04 | **9.89***, 1.27** |
| 5 | Left Somatosensory | 1 | -54 | -24 | 36 | 6160 | 2.80 | 1.48 | 0.15 | **8.56***, 1.39** |
| 6 | Cerebellum | 1 | -38 | -46 | -40 | 1720 | 0.20 | 3.60 | 2.61 | **8.84***, 1.14** |
| 7 | Left Posterior Cingulate | 1 | -12 | -26 | 40 | 3536 | 0.06 | 0.74 | 0.00 | **10.09***, 1.1** |
| 8 | Cerebellum | 1 | 14 | -56 | -54 | 3536 | 1.03 | 1.34 | 0.30 | **8.61***, 1.09** |
| 9 | Left Superior Parietal | 1 | -24 | -60 | 58 | 36728 | 0.93 | 1.21 | **5.53*** | **11.04***, 2.02** |
| 10 | Right Posterior Cingulate | 1 | 14 | -22 | 42 | 1312 | 0.65 | 1.34 | 0.07 | **8.57***, 1.15** |
| 11 | Brainstem/Medulla | 1 | -8 | -22 | -42 | 1824 | 3.04 | **6.90*** | 0.07 | **7.52***, 1.16** |
| 12 | Brainstem/Pons | 1 | -10 | -32 | -8 | 2352 | 1.81 | 3.19 | 2.89 | **9.77***, 1.14** |
| 13 | Left Middle Temporal Cortex | 2 | -52 | -26 | -12 | 4624 | 0.13 | 0.20 | 0.21 | **-7.40***, 1.32** |
| 13 | Left parahippocampal/ fusiform | 1 | -26 | -40 | -20 | 1600 | **6.14*** | 3.69 | **31.35*** ** | **5.93***, 0.84** |
| 14 | Right parahippocampal/ fusiform | 1 | 30 | -40 | -18 | 416 | 3.14 | 0.22 | **18.89*** ** | **5.14***, 0.66** |
| 15 | Right DLPFC | 1 | 32 | 38 | 34 | 840 | 0.49 | 2.40 | **15.61*** ** | **4.79***, 0.94** |
| 16 | Left Putamen | 1 | -22 | 4 | 4 | 600 | 0.003 | 1.87 | **6.06*** | **6.26***, 0.87** |

| | | | | | | | | | | |
|---|---|---|---|---|---|---|---|---|---|---|
| 17 | Right Posterior Insula | 1 | 40 | -28 | 16 | 536 | 2.48 | 0.43 | 1.42 | **7.12***, 1.02** |
| 18 | Right TPJ | 1 | 64 | -30 | 22 | 904 | 3.03 | 2.71 | 0.85 | **7.60***, 0.89** |
| 19 | Left Motor | 1 | -54 | 2 | 34 | 5288 | 0.14 | 2.21 | 0.98 | **9.23***, 1.44** |
| 20 | Left Middle Frontal Gyrus | 2 | -46 | 14 | 46 | 4872 | 0.43 | 0.00 | 0.34 | **-7.29***, 1.42** |
| 21 | Inferior Parietal Lobule | 2 | -50 | -56 | 54 | 11432 | 1.72 | 1.28 | 0.10 | **-8.53***, 1.44** |
| 23 | Left Temporal Pole | 2 | -48 | 0 | -44 | 1312 | 1.93 | 0.01 | 0.92 | **-5.85***, 1.10** |
| 24 | PFC | 2 | -6 | 28 | 60 | 142512 | 0.04 | 0.86 | 0.35 | **-14.08***, 2.43** |
| 24a | dmPFC[a] | 2 | 10 | 46 | 50 | 21648 | 0.0018 | 0.0239 | 0.0005 | **-15.00***, 2.81** |
| 24b | Left vlPFC[a] | 2 | -42 | 22 | -10 | 7576 | 0.000 | 0.0154 | 0.0057 | **-9.79***, 2.35** |
| 24c | Right vlPFC[a] | 2 | 44 | 22 | -14 | 8736 | 0.0018 | 0.0212 | 0.0112 | **-10.89***, 2.19** |
| 24d | Left dlPFC[a] | 2 | -22 | 50 | 26 | 5128 | 0.0011 | 0.0000 | 0.0029 | **-15.18***, 2.26** |
| 24e | Right dlPFC[a] | 2 | 46 | 16 | 50 | 656 | 0.0001 | 0.0010 | 0.0101 | **-7.52***, 1.46** |
| 25 | Angular Gyrus | 2 | 46 | -62 | 52 | 10632 | 1.42 | 1.19 | **5.08*** | **-8.58***, 1.46** |
| 26 | Cerebellum | 2 | -38 | -80 | -38 | 26392 | 0.01 | 0.68 | 0.43 | **-8.83***, 1.42** |
| 27 | Right Middle Temporal Cortex | 2 | 62 | -26 | -18 | 6656 | 0.08 | 1.40 | 0.09 | **-12.00***, 1.56** |
| 28 | Accumbens/sgACC | 3 | -4 | 24 | -4 | 800 | 0.004 | **30.16*** | **8.38** | **5.49***, 1.06** |
| 29 | Right Anterior Insula | 3 | 46 | -6 | -8 | 928 | 2.85 | 0.66 | 2.76 | **8.56***, 1.11** |
| 30 | Posterior Cingulate | 3 | 2 | -26 | 26 | 3832 | 0.21 | 1.18 | 1.58 | **-6.80***, 1.34** |
| 31 | Visual Cortex | 3 | 10 | -74 | 28 | 38424 | 0.00 | 0.44 | 0.31 | **-8.77***, 1.30** |

* p < 0.05, ** p < 0.01, *** p<0.001

a. The prefrontal cortex brain region was divided into smaller subclusters using a higher threshold in the table, and tested for video effects in the follow up analyses. These subclusters were not used for the cluster-level analysis.
